# High innate preference for black substrate in the chive gnat, *Bradysia odoriphaga* (Diptera: Sciaridae)

**DOI:** 10.1101/511121

**Authors:** Lina An, Fan Fan, Klaus Lunau, Mengyao Li, Xiaofan Yang, Guoshu Wei

**Author notes:** Corresponding author: Prof. Dr. Guoshu Wei, College of Plant Protection, Hebei Agricultural University.

## Abstract

The chive gnat, *Bradysia odoriphaga*, is a notorious pest of *Allium* species in China. Colour trapping is an established method for monitoring and controlling of *Bradysia* species. In order to clarify the effect of colour preference of *B. odoriphaga* for the egg-laying substrate, multiple-choice tests were employed to assess the spontaneous response of the chive gnat to different colour hues and brightness levels under different intensities of white illumination and two spectrally different illuminations. Given the choice among four colours differing in hue under different intensities of white illumination and two spectrally different illuminations, chive gnat adults visited preferably the black substrate, a lesser extent to brown and green substrates, and the least extent to orange substrate irrespective of illumination. Given the choice among four levels of brightness under the same illumination conditions as those in the previous experiment (different intensities of white illumination and two spectrally different illuminations), chive gnats preferred black substrate over dark grey, and these over light grey and white substrates. Meanwhile, both virgin and copulated adults significantly preferred black over other colour hues and brightness. Based on our results, we conclude that the chive gnat adults significantly prefer black substrates irrespective of colour hues and brightness. This behaviour does not alter due to ambient light condition changes. No difference observed between choices of female and male adults. Our results provide new insight for understanding the colour choice behaviour in chive gnat and pave a way to improve monitoring and control of chive gnats and management.

**Summary statement:** Chive gnat (*Bradysia odoriphaga*) innately prefer to move to black substrate irrespective of colour hues and brightness. This behaviour maintained the ambient lights change.

## Introduction

The chive gnat, *Bradysia odoriphaga* (Diptera: Sciaridae), is the most destructive pest to *Allium* vegetables in China, especially to Chinese chive *Allium tuberosum*. Although chive gnat adults do not cause plant damage, the females lay eggs around the root in soil, hatch into larvae that directly damage roots and bulbs of plants, thus disrupting the uptake of water and nutrients (Mei et al., 2003). Historically, the control of chive gnat has been dependent on the use of chemicals, such as chlorpyrifos and phoxim (Gao et al., 2000; Mu et al., 2005), but it didn`t turn out well mainly due to the cryptic larval life style and the development of resistance to insecticides (Zhang et al., 2003). Particularly, excessive use of certain pesticides will lead to environmental pollution and high pesticide residues. Therefore, it`s necessary and exigent to search safe and efficient management strategies to control chive gnat.

Many insects use visual stimuli to perceive a variety of resources, such as adult food, mating encounter sites, oviposition sites or shelter from harmful biotic or abiotic conditions (Labeyrie, 1978; Southwood, 1973). The quality of the perception of visual objects, however, is strongly influenced by the characteristics of reflected light including hue and brightness. Colour is especially important for distinguishing resource quality, e.g. flower condition, partner selection (von Frisch 1914; Kelber 2006; An et al., 2018; Koethe et al., 2018), as well as location (e.g. oviposition site, shelter) (Osorio and Vorobyev, 2008; Collins and Blackwell, 2000). In this study we investigated the preference of chive gnat for colour and brightness of chive gnat in order to reduce the damage of *A. tuberosum* by means of monitoring or controlling the chive gnat.

Vision-orientated coloured sticky traps may represent relevant potential monitoring and control strategies of *Bradysia* species (Cloyd et al., 2007), since these trapping methods are environment-friendly and do not cause pesticide residues and pesticide resistance. Colour trapping is a common method for trapping various insect species (Gao et at., 2016). Many insects have already been confirmed to exhibit colour preferences including those for distinct colour hues, colour saturation, colour brightness, and colour contrast (Lunau and Maier, 1995; Chittka and Menzel, 1992; but see Kelber, 2005). Most profound studies about innate colour preferences in insects focus on pollinating insects such as bees (Lunau et al., 1996; Hempel de Ibarra et al., 2000; Koethe et al., 2018), lepidopterans (Weiss, 1997; Goyret et al., 2008), and flies (Ilse, 1949; An et al. 2018; Lunau et al., 2018), whereas studies about colour preferences in agricultural pests mostly evaluate the results of colour trapping (Bian et al., 2016; Silva et al., 2018). Colour trapping is a common method for the control of dipteran pest species. For example, the whitefly, *Bemisia tabaci*, some tephritid flies, and anthomyid flies, *Strobilomyia* spp. are particularly attracted by yellow sticky traps (Hill and Hooper, 2011; Jenkins and Roques 1993; Hou et al., 2006), whereas fungus gnat, *Bradysia difformis*, exhibit an innate colour preference for black (Stukenberg et al., 2018). Actually, in field experiments with coloured sticky traps chive gnat have already been successfully captured (Hong, 2016), but the quantitative analysis of the contribution of colour parameters such as hue and brightness to lure chive gnats has never been concerned so far.

The purpose of our experiments was to determine the relative attractiveness of different colours to adult chive gnats and to assess the efficacy of colour parameters, hue and brightness, to attract chive gnats. In addition we investigated whether the chive gnat adults maintained their innate colour preference when the colour stimuli were presented under various light intensities of white illumination and two spectrally different illuminations. The outcome of these experiments will lead to a better understanding of their colour choice behaviour and colour vision and is thereby beneficial to understand their biological characteristics and develop specific monitoring tools and efficient control strategies.

## Material and methods

### Chive gnat rearing and handling

*B. odoriphaga* larvae were initially obtained from a field of *Allium tuberosum* in Cangzhou Hebei Province, China during May 2018. The colonies of *B. odoriphaga* were maintained in the IPM Laboratory of Hebei Agricultural University and reared on *A. tuberosum* for more than 6 generations. Eggs, larvae and pupae were reared in Petri dishes (9cm in diameter, 2.5cm height) containing a filter paper that was soaked with 2.5% agar medium, and the fresh chive *A. tuberosum* was placed in a separate petri dish as diet for the larvae. The chive gnats were placed in rearing containers made of plastic pots (9cm top diameter, 15cm bottom diameter, 5cm height). A petri dish (15cm diameter) was used as the bottom of each rearing container, and a reversed plastic cup (9cm top diameter, 15cm bottom diameter, 5cm height; with dozens of needle holes for gas exchange) was used as the cover. The container between the bottom and the cover was sealed with sealing film. Each petri dish contained a filter paper that was soaked with 2.5% agar medium (about 15ml) for maintaining moisture. Newly emerged female and male gnats could mate immediately. Female gnats can lay eggs one or two days after mating. Insect colonies were maintained in climate chambers maintained at 24±1°C with 75±5% relative humidity and a 14:10 hours light: dark cycle.

### Colour hue and brightness

Based on the colours of chive gnats` body surface, of the fresh and old host plants, and of the soil green, orange, brown, and black colour papers were selected as stimuli for the experiment with varying colour hues. Four brightness levels including white, light grey, dark grey, and black and two blue colour stimuli differing in brightness were selected for the experiment with varying colour brightness. Colour papers made of photographic paper printed via POWERPOINT printed by a colour inkjet printer (HP 100) were offered (Table 1). The spectral reflectance of the colour stimuli was measured by a spectrophotometer (Konica Minolta CM-3700A, Japan) (Fig. 1). Light emitting diodes used in our experiment were designed specific ranges of wavelengths, such as that of green light from 525 to 530nm, of blue light from 455 to 460nm and white light with a colour temperature of 6000~6500K. All intensities of LEDs were measured by illuminometer (TES-1339, Tes Electronics Industry Corporation, China)

**Table 1.** The RGB values of the different colour hues and brightness levels.

| Experiment | Colour stimuli | RGB value |
| --- | --- | --- |
| Colour hues | Black | 0, 0, 0 |
|  | Brown | 120, 60, 0 |
|  | Green | 0, 145, 65 |
|  | Orange | 245, 140, 0 |
| Brightness levels | Black | 0, 0, 0 |
|  | White | 255, 255, 255 |
|  | L-grey | 190, 190, 190 |
|  | D-grey | 90, 90, 90 |

**Fig. 1.**
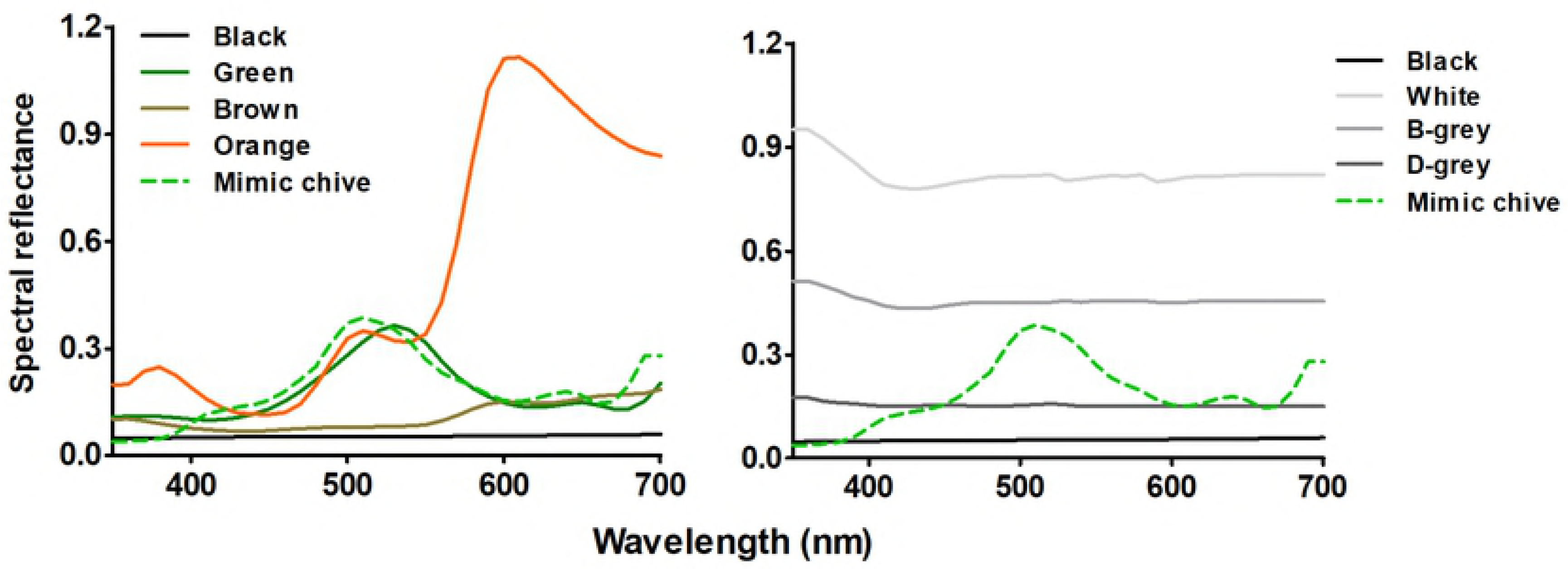
Reflectance spectra of all colour stimuli used in the experiments. L-grey means light grey, D-grey means dark grey, L-blue means light blue, D-grey means dark grey.

### Experimental device

The device for multiple choice tests is a quadrilateral cube (length×width× height=30×30×30 cm) made out of cardboards and has four chambers of identical size. In the middle of the device is the release zone displaying a white colour (length×width=8×8 cm) of flies to be tested (Fig. 2). Each chamber contained an artificial Chinese chive which was placed in the middle of the chambers` bottom. The device has a lid made of Plexiglas, which was used to prevent chive gnat adults from flying out of the device (Fig. 2).

**Fig. 2.**
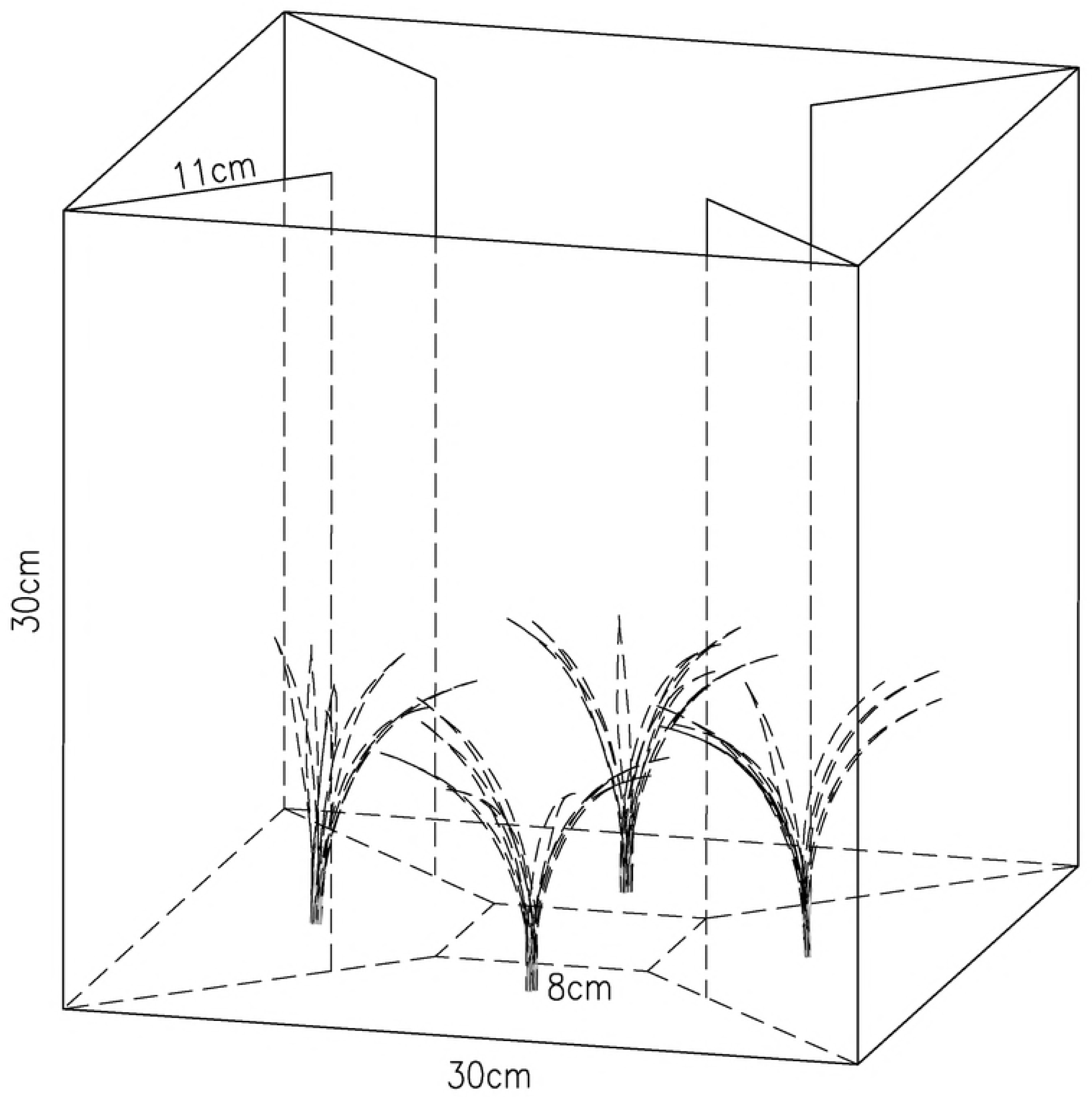
Device of the colour choice tests with *Bradysia odoriphaga.* Scheme of the device of colour choice test, which was designed as a cube (edge length=30cm). In the middle of cube was small square (edge length=8cm) used as a decision area. The chambers were separated by cardboard. All four differently coloured chambers contained a mimic chive plant.

### Experimental procedure

Based on the activity characteristics of chive gnat adults (Hong et al., 2017), all the experiments were started at 9:00 am every day. The experimental conditions simulated those of greenhouses used for growing Chinese chive. Before the tests the chive gnat adults were placed in a completely dark environment for 30 min for equal adaptation. In the experiments of innate preference for colour hues and brightness levels, each chamber was pasted with one of four coloured papers to be used as a colour chamber for quadruple choice.

### Experiment 1: Colour hue and brightness preference under four intensities of white illumination

The rationale of the experiments was to study the innate preference of chive gnat adults to respond different colour hues and brightness of stimuli, respectively. Four colours differing in hues, black, orange, brown and green, were used to test the colour preference of chive gnats for four different intensities of ambient light. In addition, four brightness levels of stimuli, white, light grey, dark grey and black, were used to test the brightness preference of chive gnat for four different intensities of ambient light. For each trial 30 newly emerged, healthy adults were put into the release zone of the device under white light with 0.1, 100, 1000, 10000lux, respectively. And the device was immediately covered with a transparent lid. After 30min the number of flies in each chamber was counted. Each treatment was repeated 20 times. Ten trials were performed with females and 10 trials were performed with males.

### Experiment 2: Colour and brightness discrimination under two spectrally different illuminations

The rationale of the experiments was to study whether chive gnat adults can maintain the colour preference when tested under spectrally different illuminations. The same colour stimuli, four colour hues and four brightness, were respective used to test the colour choice of chive gnats under blue or green illumination with 250lux. For each trial 30 newly emerged, healthy adults were put into the release zone of the device under blue and green light with 500lux, respectively. And the device was immediately covered with a transparent lid. After 30min the number of flies in each chamber was counted. Each treatment was repeated 20 times. Ten trials were performed with females and 10 trials were performed with males.

### Experiment 3: Colour hues and brightness of chive gnat adults with different physiological state

The rationale of the experiments was to study the innate preference of virgin and copulated adults to respond different colour hues and brightness of stimuli, respectively. For each trial, virgin adults and copulated adults were put into the release zone of the device under white light with 100lux, respectively. And the device was immediately covered with a transparent lid. After 30min the number of flies in each chamber was counted. Each treatment was repeated 20 times. Ten trials were performed with females and 10 trials were performed with males.

## Results

### Experiment 1: Colour and brightness preference under four intensities of white illumination

In the colour preference experiments the adults significantly preferred the black colour irrespective of light intensities while other colours, i.e. green, brown and orange, were less attractive (Fig. 3). The preference for black increased with the intensities of light increased, except for the test with 1000lux intensity. They visited the black with a choice frequency of 40.26% for 0.1lux, 47.54% for 100lux, 45.37% for 1000lux and 49.66% for 10000lux intensity, respectively. The brown and green colours were significantly less attractive. The orange colour was seemingly the least attractive with a choice frequency of 13.84% for 0.1lux intensity, 11.19% for 100lux intensity, 11.61% for 1000lux intensity and 14.78% for 10000lux intensity, respectively.

**Fig. 3.**
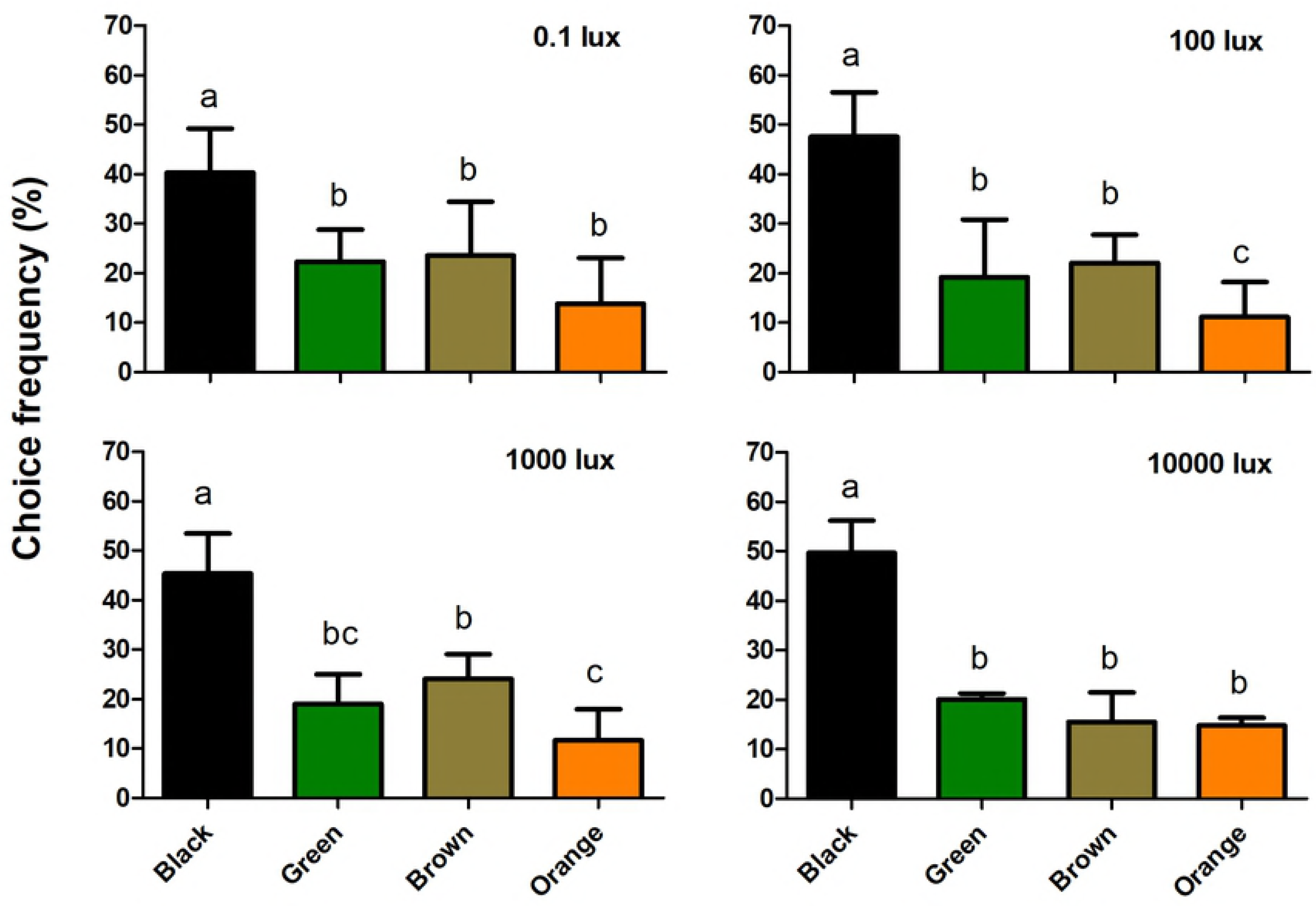
Colour choices in chive gnat *Bradysia odoriphaga* adults of different colour hues under four different intensity of white illumination. Different letters refer to significant differences according to One-way ANOVA with P<0.05.

In the brightness choice experiments the adults also significantly preferred the black colour irrespective of light intensities, while other brightness levels were less attractive (Fig. 4). They visited the black with a choice frequency of 36.73% for 0.1lux, 48.72% for 100lux, 44.39% for 1000lux and 49.75% for 10000lux intensity, respectively. The choice frequency for dark grey was 28.47%, 23.65%, 26.01% and 30.52%, respectively. The light grey and white colour were less attractive. The chive gnats visited the light grey and white with a choice frequency of 20.12% (light grey) and 14.67% (white colour) for 0.1lux; 16.11% (light grey) and 11.56% (white colour) for 100lux; 19.06% (light grey) and 10.54% (white colour) for 1000lux and 11.23% (light grey) and 8.5% (white colour) for 10000lux intensity. The control experiment using the device with the same colour (white colour) in each chamber under 100 lux white illumination showed that chive gnats did not prefer one of the chambers (supplement S1). The colour preference was similar for males and females (Table 2).

**Fig. 4.**
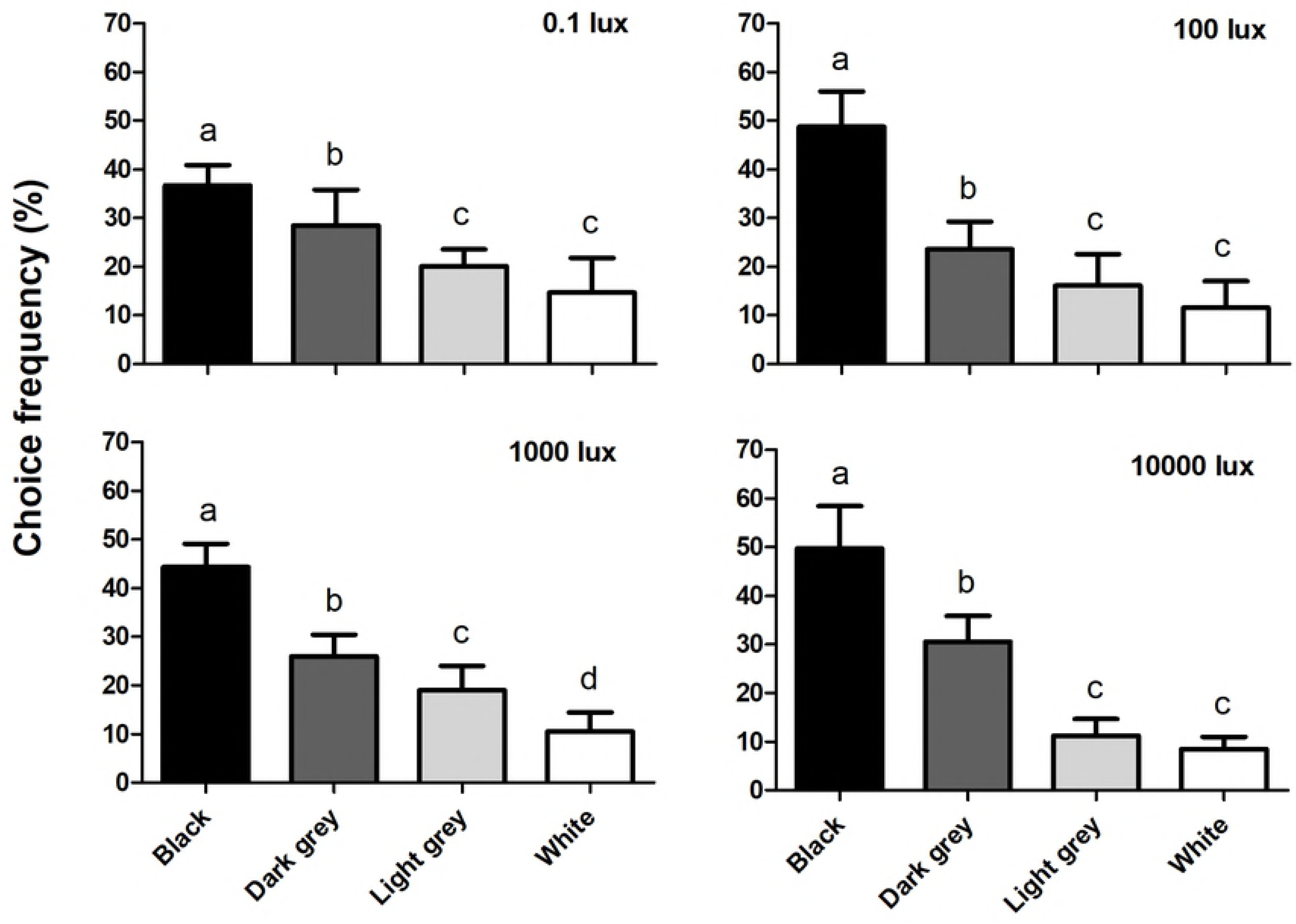
Colour choices of stimuli varying in brightness in chive gnat *Bradysia odoriphaga* adults under four different intensities of white illumination. Different letters refer to significant differences according to One-way ANOVA with P<0.05.

**Table 2.** Variance analysis of the response of *Bradysia odoriphaga* to colour hues and brightness under different light

| Experiment | Influencing factors | df | Mean square | F | Sig. |
| --- | --- | --- | --- | --- | --- |
| exp.1<br>Colour hues<br>and<br>Brightness | Colour hues | 3 | 4928.634 | 237.479 | 0.000 |
|  | Sex | 1 | 4.167E-006 | 0.000 | 1.000 |
|  | Light intensity | 3 | 2.639E-005 | 0.000 | 1.000 |
|  | Colour hues * Light intensity | 9 | 96.965 | 4.672 | 0.000 |
|  | Colour hues * Sex | 3 | 14.646 | 0.706 | 0.552 |
|  | Light intensity * Sex | 3 | 4.167E-006 | 0.000 | 1.000 |
|  | Colour hues * Light intensity * Sex | 9 | 41.156 | 1.983 | 0.056 |
|  | Brightness | 3 | 5003.351 | 243.417 | 0.000 |
|  | Sex | 1 | 5.104E-005 | 0.000 | 0.999 |
|  | Light intensity | 3 | 2.326E-005 | 0.000 | 1.000 |
|  | Brightness * Light intensity | 9 | 131.531 | 6.399 | 0.000 |
|  | Brightness * Sex | 3 | 66.062 | 3.214 | 0.029 |
|  | Light intensity * Sex | 3 | 2.326E-005 | 0.000 | 1.000 |
|  | Brightness * Light intensity * Sex | 9 | 10.308 | 0.502 | 0.868 |
| exp.2<br>Colour hues<br>and<br>Brightness | Colour hues | 3 | 2176.384 | 22.634 | 0.000 |
|  | Sex | 1 | 2.083E-006 | 0.000 | 1.000 |
|  | Wavelength | 1 | 2.083E-006 | 0.000 | 1.000 |
|  | Colour hues * Sex | 3 | 160.182 | 1.666 | 0.194 |
|  | Colour hues * Wavelength | 3 | 26.082 | 0.271 | 0.846 |
|  | Wavelength * Sex | 1 | 2.083E-006 | 0.000 | 1.000 |
|  | Colour hues * Sex * Wavelength | 3 | 98.290 | 1.022 | 0.396 |
|  | Brightness | 3 | 2449.824 | 41.924 | 0.000 |
|  | Sex | 1 | 8.333E-006 | 0.000 | 1.000 |
|  | Wavelength | 1 | 8.333E-006 | 0.000 | 1.000 |
|  | Brightness * Sex | 3 | 58.165 | 0.996 | 0.407 |
|  | Brightness * Wavelength | 3 | 17.611 | 0.302 | 0.824 |
|  | Wavelength * Sex | 1 | 0.000 | 0.000 | 1.000 |
|  | Brightness * Sex * Wavelength | 3 | 1.748 | 0.030 | 0.993 |
| exp.3<br>Colour hues<br>and<br>Brightness | Colour hues | 3 | 97182.366 | 22111.831 | 0.000 |
|  | Sex | 1 | 25.843 | 5.880 | 0.508 |
|  | Physiological state | 1 | 0.689 | 0.157 | 0.692 |
|  | Colour hues * Sex | 3 | 145.064 | 33.006 | 0.000 |
|  | Colour hues * Physiological state | 3 | 26.087 | 5.936 | 0.001 |
|  | Physiological state * Sex | 1 | 0.707 | 0.161 | 0.688 |
|  | Colour hues * Sex * Physiological state | 3 | 29.337 | 6.675 | 0.000 |
|  | Brightness | 3 | 97746.048 | 28301.236 | 0.000 |
|  | Sex | 1 | 0.265 | 0.077 | 0.782 |
|  | Physiological state | 1 | 8.887 | 2.573 | 0.109 |
|  | Brightness * Sex | 3 | 190.087 | 55.038 | 0.000 |
|  | Brightness * Physiological state | 3 | 250.019 | 72.390 | 0.000 |
|  | Physiological state * Sex | 1 | 32.972 | 9.547 | 0.002 |
|  | Brightness * Sex * Physiological state | 3 | 149.622 | 43.321 | 0.000 |

Note: All the data were analyzed by Multifactor Variance Analysis (SPSS 17.0)

### Experiment 2: Colour and brightness discrimination under two spectrally different illuminations

Given the multiple choice among four colours differing in hues, black, blue, brown and orange, with blue light of 500lux intensity, the choice frequency of chive gnats for black was 42.77%, which was significantly higher than those for blue, brown and orange colours (19.67%, 22.09% and 15.47%). A similar result was obtained testing the chive gnats in green light of 500lux intensity; 44.45% of the adults significantly preferred the black than other three colours with a choice frequency of 16.83%, 25.49% and 13.22%, respectively (Fig. 5A).

**Fig. 5.**
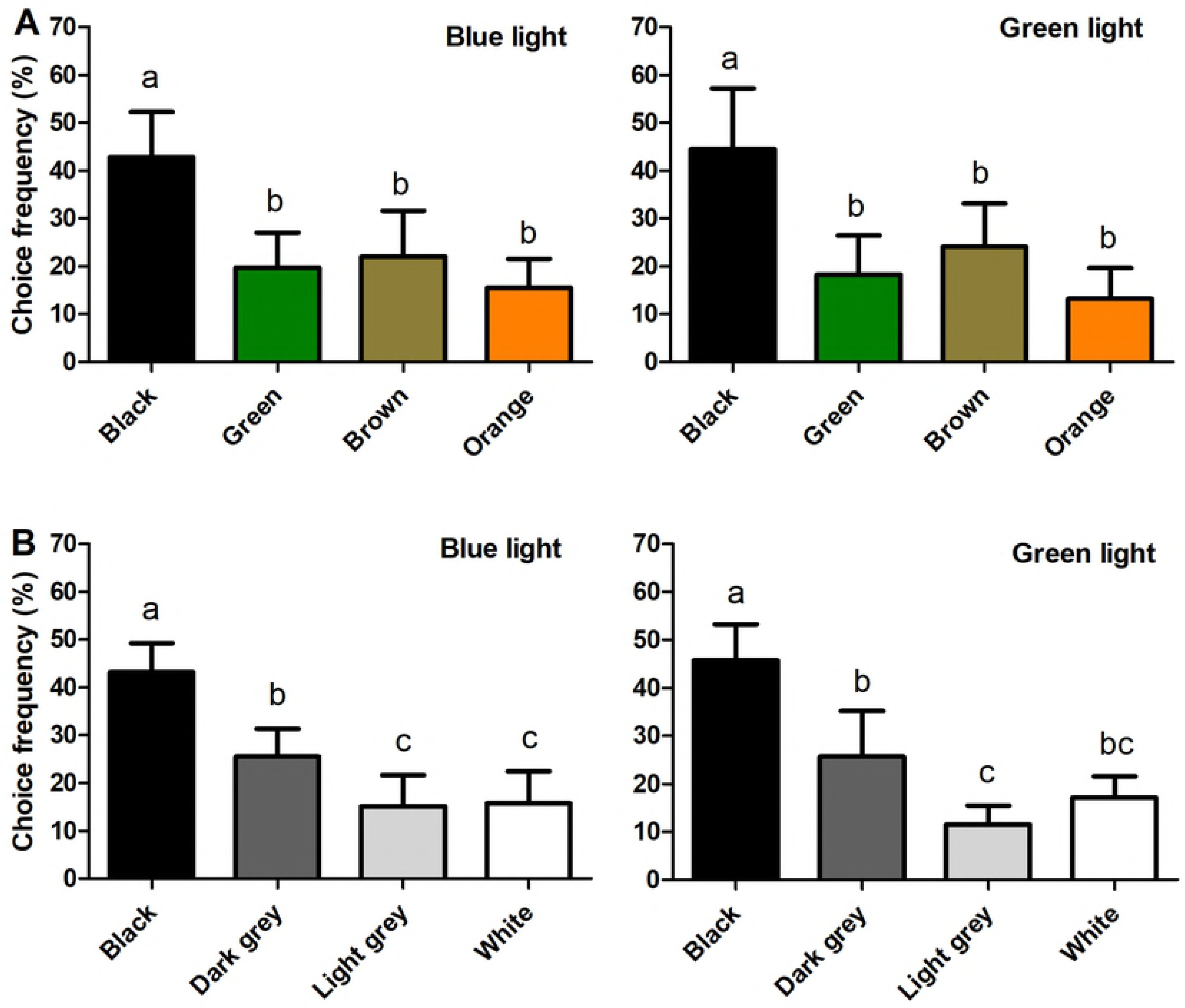
Colour and brightness preference of chive gnat *Bradysia odoriphaga* adults under two spectrally different illuminations. (A) Colour choice among four colour stimuli differing in hue under blue light (left) and green light (right). (B) Choice among four colour stimuli differing in brightness under blue light (left) and green light (right). Different letters refer to significant differences according to One-way ANOVA with P<0.05.

Moreover, the multiple choice among four levels of brightness, black, dark grey, light grey and white, under blue light of 500lux intensity were used for testing the choice preference of the chive gnat adults. The adult chive gnats significantly preferred black with a percentage of choice amounting to 43.16% over dark grey amounting to 25.55% as well as light grey and white amounting to 18.88% and 12.41%, respectively. Also under green light of 500lux intensity the chive gnat adults significantly preferred black with a choice frequency amounting to 45.73% over dark grey, light grey and white amounting to 25.64%, 18.23% and 10.40%, respectively (Fig. 5B).

### Experiment 3: Colour hues and brightness of chive gnat adults with different physiological state

In the colour choice experiments both the virgin and copulated adults significantly preferred the black colour, to a lesser visited brown and green, whereas orange was significantly less attractive (Fig. 6). By contrast, in the brightness experiment, both the virgin and copulated adults significantly preferred the black colour over dark grey, and those over light grey and white (Fig. 6).

**Fig. 6.**
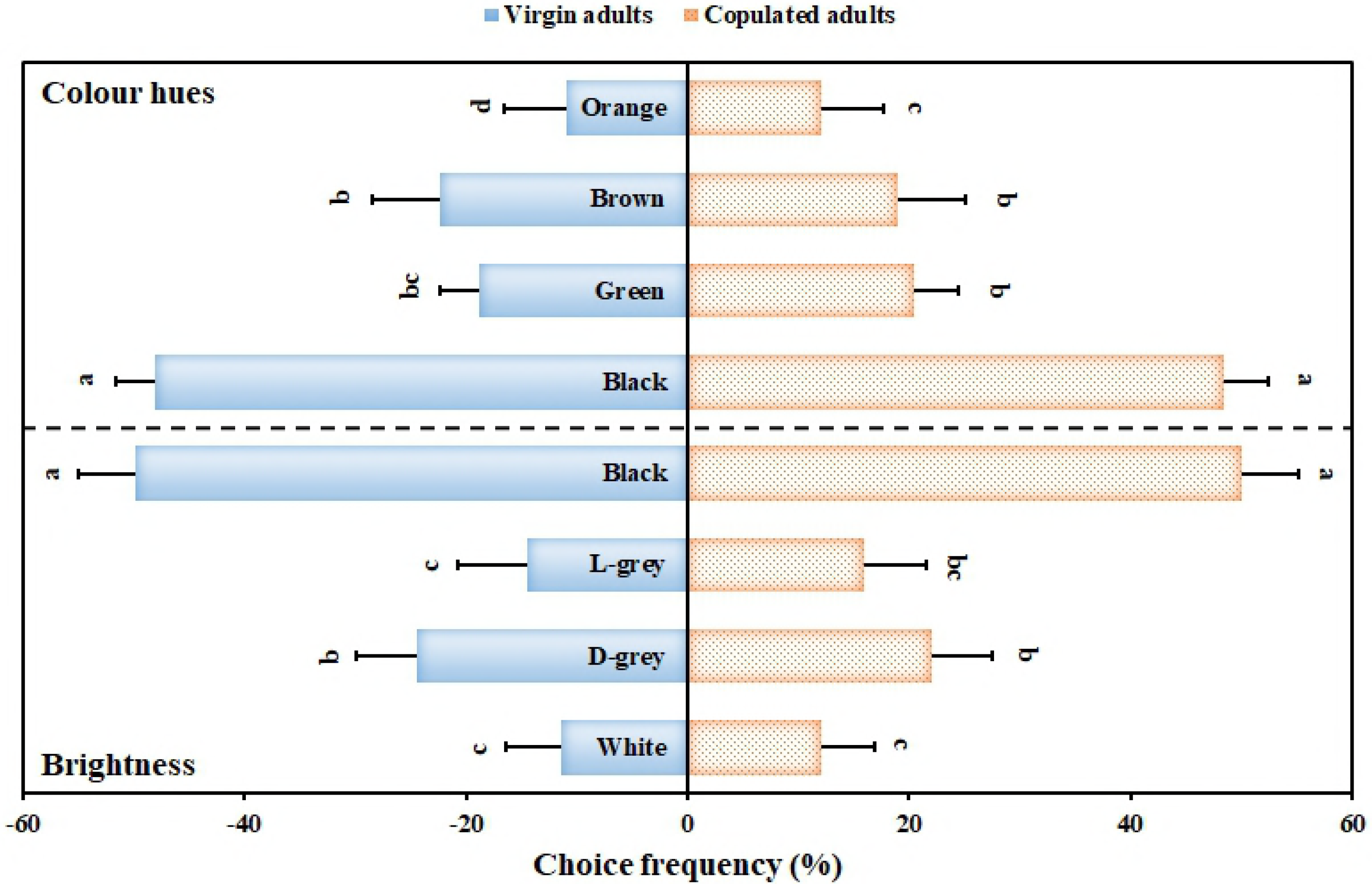
Colour and brightness preference of chive gnat *Bradysia odoriphaga* adults with different physiological states Different letters refer to significant differences according to One-way ANOVA with P<0.05.

In all experiments there was no significant difference in colour choice behaviour between female and male in chive gnat, and also no significant differences in colour preference between virgin and copulated adults were found (Table 2).

## Discussion

Adult chive gnats, *Bradysia odoriphaga*, showed a significant preference for the black substrate, while other coloured substrates attracted only a limited number of chive gnat. These results provide strong evidence that chive gnats possess an innate colour preference for black substrates and some evidence that they maintain the preference for black even if the ambient light conditions change, i.e. the preference for black is not altered by intensity and spectral composition of the illuminating light. Although the tests were specifically designed to capture female chive gnats referring to the egg-laying sites, no differences in the colour preference between the virgin and copulated adults of *B. odoriphaga* were found and there were no differences in the colour preference between female and male. We, therefore, speculated chive gnat adults prefer black substrate not only for searching oviposition sites, but also for other reasons, such as searching for mates or finding a safe place for hiding in camouflage due to their black surface. Colour traps have been used to control *Bradysia* gnats under field conditions (Ma et al., 2013; Hong, 2016), but innate colour preferences have never been investigated in detail for chive gnats, *B. odoriphaga*.

Wavelength-specific behaviours known from specific tasks such as oviposition in butterflies (Lepidoptera) and flies (Diptera) regularly are dependent of intensity (Song and Lee, 2018). Gravid females of certain mosquitos, *Toxorhynchites moctezuma* and *T. amboinensis* (Collins and Blackwell, 2000), oviposited preferentially into black substrate. Remarkably, the flower-visiting hoverfly *Eristalis tenax* prefers yellow colours, can learn many other colours, but strongly avoids dark colours (An et al., 2018). The canopy ant, *Cephalotes atratus*, prefers bright white colours when given a choice of target colours of varying shades of grey; specifically brightness seems to have a great influence on the landing behavior of canopy ants, thus it is suspected that the high contrast between tree trunks and the darker surrounding foliage provides the preferred visual target for falling ant (Yanoviak and Dudley, 2006).

Likewise other dipteran pests are known to be attracted to black colours such as the bluebottle fly, *Calliphora vomitoria*, (Benelli et al., 2018) and the tabanid fly, *Tabanus illotus*, (Bracken et al., 1962). The preference for black surfaces found in water-living insects (Schwind, 1995) and tabanid flies (Horwarth et al., 2008) is associated with the perception of horizontally polarized light reflected from shiny surfaces such as water which is optimally seen at black targets (Horwarth et al., 2008). Since the target colours used in the colour choice tests with the chive gnat were not shiny and the light source was not the sun, it is very unlikely that polarization vision might have influenced the colour choice of the chive gnat.

The black surface was preferred by chive gnats in comparison to all other colours. One possible reason is that the chive gnats performed a colourblind choice relying only on the contrast between the black target and other colours and the background which is one of the key features for object perception of insects (Prokopy and Owens, 1983) including Diptera (Lunau, 2014). A strong brightness contrast may be found in nature between light plant stems, leaves and the dark substrate. The finding that adult fungus gnats, *Bradysia difformis,* significantly preferred black sticky traps over yellow ones, was interpreted that fungus gnat adults were searching for a convenient egg-laying substrate (Stukenberg et al., 2018). Based on this hypothesis, the spectral reflectance of the host plant (*Allium tuberosum*) and soil substrate close to root of the chive gnat was measured (Fig. 7). The maximum value of spectral reflectance of leaves (*A. tuberosum*) is 38%, whereas the maximal reflectance of soil substrate is about 9%, which is very close to the value of the black colour in our experiments (black: 6%). As a conclusion, we assume that the strong colour contrast between the substrate for egg-laying and the host plant might guide the search for mating partners or oviposition sites in dim surroundings.

**Fig. 7.**
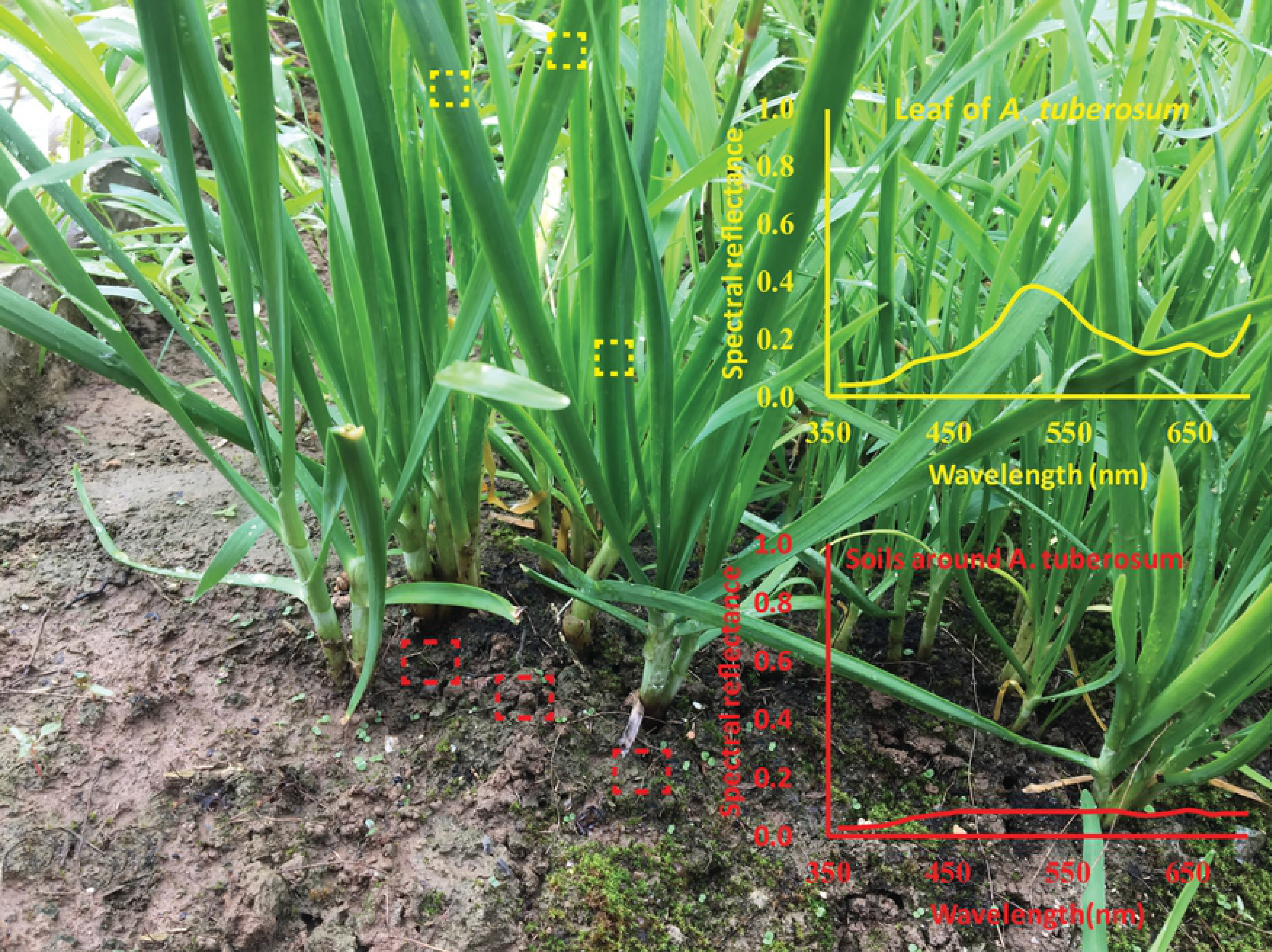
Reflectance spectra of living environment in chive gnat *Bradysia odoriphaga*. Dashed boxes with yellow colour represent the measure sites of leaves of *A. tuberosum*. Dashed boxes with red colour represent the measure sites of soils around *A. tuberosum*.

Although habitat-related olfactory stimuli have been identified as important cues in the specific task of some insects (Riffell 2012; Clifford and Riffell 2013; Linley 1988, 1989), visual stimuli should not be overlooked as an important sensory modality, especially the aspect of searching and finding host plant (Reeves, 2011). In our study, we focused on the effect of visual stimuli for chive gnats, not olfactory stimuli, so a scentless artificial host plant was used as an attractive signal in order to avoid the odour interference of host plant. Further studies are needed to explore the interaction of visual and olfactory cues for oviposition and the visual mechanism underlying the colour choice in chive gnats, i.e. photoreceptor types, visual system and their visual ecological significance.

